# Transcriptionally active nitrogen fixation and biosynthesis of diverse secondary metabolites by *Dolichospermum* and *Aphanizominom*-like Cyanobacteria in western Lake Erie Microcystis blooms

**DOI:** 10.1101/2022.09.30.510322

**Authors:** Colleen E. Yancey, Olivia Mathiesen, Gregory J. Dick

## Abstract

Cyanobacterial harmful algal blooms (cyanoHABs) in the western basin of Lake Erie are dominated by microcystin producing *Microcystis* spp., but other cyanobacterial taxa that coexist in these communities may play important roles in production of toxins and shaping bloom dynamics and community function. In this study, we used metagenomic and metatranscriptomic data from the 2014 western Lake Erie cyanoHAB to explore the genetic diversity and biosynthetic potential of cyanobacteria belonging to the *Anabaena, Dolichospermum, Aphanizomenon* (ADA) clade. We reconstructed two near-complete metagenome-assembled genomes from two distinct ADA clade species, each containing biosynthetic gene clusters that encode novel and known secondary metabolites that were transcriptionally active. These taxa also appear to have varying nutrient acquisition strategies, and their ability to fix N may be important for synthesizing N rich metabolites as well as supporting bloom persistence. Although not the dominant organism in this system, these results suggest that ADA may be important community members in western Lake Erie cyanoHABs that have the potential to produce unmonitored toxins.

**Highlights:**

- Through metagenomic approaches, we generated two near-complete metagenome assembled genomes from two distinct species that are dispersed across the ADA clade of cyanobacteria.
- These ADA cyanobacteria have the potential to produce a variety of known and novel secondary metabolites, and use different nitrogen fixation strategies as observed through differential transcript abundance
- This works highlights the diversity of cyanobacteria in western Lake Erie blooms despite their continued dominance by *Microcystis*, and that these less abundant cyanobacteria may produce unmonitored toxins and shape bloom dynamics through N-fixation.

## 1. Introduction

For the past 20 years, western Lake Erie has endured annual cyanobacteria Harmful Algal Blooms (cyanoHABs) (Bridgeman et al., 2013; Stumpf et al., 2012) dominated by *Microcystis* spp. (Berry et al., 2017; Bridgeman et al., 2013; Rinta-Kanto et al., 2005, 2009). *Microcystis* spp. are believed to have gained dominance in this system via excessive N-loading (Davis et al., 2010; Gobler et al., 2016), which enabled non-diazotrophic cyanobacteria to be ecologically successful. These contemporary blooms contrast those observed during the 1960s and 1970s, which were more diverse and dominated by diazotrophs (Davis, 1964; Makarewicz, 1993; Munawar and Munawar, 2011; Paerl et al., 2018). CyanoHABs in this system were largely reduced by the 1980s due to the Great Lakes Water Quality Agreement that implemented reductions on phosphorus (P) inputs (Matisoff and Ciborowski, 2005), as it was initially identified as the major driver of cyanoHABs (Schindler, 1974), while nitrogen (N) inputs were left unmanaged. Eventually this led to N and P co-limitation (Paerl et al., 2016; Paerl and Scott, 2010), which shifted species composition, and contributed in part to the eventual reemergence of cyanoHABs in the western basin (Watson et al., 2016).

While *Microcystis* dominates western Lake Erie blooms and has been mainly responsible for the production of the hepatotoxin microcystin, bloom communities contain other cyanobacteria including *Cyanobium* spp., *Pseudanabaena* spp., *Planktothrix* spp. (near river mouths and bays), and members of the *Anabaena, Dolichospermum, Aphanizomenon* (ADA) clade of cyanobacteria (Berry et al., 2017; Bullerjahn et al., 2016; Chaffin et al., 2013; Gobler et al., 2016). However, little is known about their role in these communities, their impact on bloom dynamics, or their potential contribution to toxin production in western Lake Erie.

The ADA clade of cyanobacteria are of particular concern as they are prolific producers of diverse, potent, cyanotoxins (Österholm et al., 2020). These toxins include microcystin (Fewer et al., 2008; Tonk et al., 2009; Vaitomaa et al., 2003), neurotoxic anatoxin (Carmichael et al., 1975), and paralytic shellfish toxin (PST) saxitoxin, which has recently been classified as a bioweapon via the Chemical Weapons Convention (Al-Tebrineh et al., 2010; Sierra and Martínez-Álvarez, 2020). While toxin production is variable and sporadic across this clade (Österholm et al., 2020), there is still the potential that these cyanobacteria can produce a variety of potent toxins within natural populations. Whereas microcystin production has largely been attributed to *Microcystis* in western Lake Erie (Berry et al., 2017; Rinta-Kanto et al., 2005, 2009; Steffen et al., 2017), and anatoxin and saxitoxin have yet to be widely detected in this system (McKindles et al., 2020), the presence of ADA species in these blooms merits further investigation given the potential for novel or emerging toxins. Indeed, ADA have dominated Lake Erie cyanoHABs in the past (Davis, 1964), saxitoxin genes have been detected in early summer *Dolichospermum* blooms in the central basin of Lake Erie (Chaffin et al., 2019), and mixing and spreading of blooms from the western to central basin have been observed (Michalak et al., 2013). A better understanding of ADA distribution, ecophysiology, and biosynthetic potential will improve our understanding of how changing nutrient regimes under various climate and nutrient management scenarios are likely to affect the abundance, distribution, and toxicity risk of these organisms.

Within western Lake Erie, ADA cyanobacteria may also play a role in cyanoHAB community dynamics via supply of nitrogen (N) and/or competition with *Microcystis.* ADA are distinct from *Microcystis* as they can fix nitrogen through differentiated heterocyst cells (Kumar et al., 2010; Wolk et al., 1994), thereby ameliorating N limitation during deplete conditions (Wood et al., 2010). However, ADA cyanobacteria share similarities with *Microcystis* as they have large biosynthetic potential and produce diverse secondary metabolites (Kehr et al., 2011; Österholm et al., 2020; Welker and von Döhren, 2006), may regulate their buoyancy through gas vesicles (Li et al., 2016; Walsby et al., 2007), and have genomes that are rich with mobile elements (Driscoll et al., 2018; Wang et al., 2012).

To explore the genetic diversity and toxin producing potential of ADA cyanobacteria more deeply in natural cyanoHAB communities, we analyzed metagenomic and metatranscriptomic datasets from the 2014 western Lake Erie cyanoHAB. *De novo* assembly was used to recover two high quality metagenome assembled genomes (MAGs) belonging to the ADA clade. From these MAGs we assessed the genetic diversity of ADA cyanobacteria in western Lake Erie, identified biosynthetic gene clusters which encode secondary metabolites, and tracked the relative expression of biosynthesis and nutrient uptake and metabolism genes.

## 2. Materials and Methods

### 2.1 Study Site and Sample Collection

Three core stations were sampled and are part of weekly cyanoHAB monitoring by the NOAA Great Lakes Environmental Research Laboratory (GLERL) (Cooperative Institute for Great Lakes Research, 2019) within western Lake Erie. Samples were collected weekly from mid-June to late October in 2014 at core stations WE2, WE4, and WE12. WE2 is close to the inlet for the Maumee River inlet (41° 45.743’N, 83° 19.874’ W), WE4 is considered an offshore site closer to the center of the basin (41° 49.595’N, 83° 11.698’W), and WE12 is near the Toledo drinking water crib (41° 42.535’N, 83° 14.989’W).

20L water samples were collected via integrated depth water sampling, in which the entire water column, from surface to 1 meter above the bottom, was continuously sampled by pump-cast. Physiochemical measurements such as pH, water temperature, and specific conductivity were measured during the research cruise. Biomass was collected by filtering 2L of integrated depth water though a 100 μm polycarbonate mesh filter. The biomass on the filter was then collected and filtered through a 0.22 μm filter and preserved in 1 mL of RNALater^™^ (Invitrogen^™^, Ambion^™^) and placed on ice.

### 2.2 DNA Extraction and Sequencing

Extraction and sequencing have been described previously in greater detail (C. E. Yancey et al., 2022). Qiagen DNeasy Blood and Tissue Kits were used to extract DNA while The Qiagen RNEasy kit was used to complete RNA extraction and cDNA library preparation (Qiagen, Hilden, Germany). Shotgun DNA and RNA sequencing was completed at the University of Michigan Sequencing Core using the Illumina^®^ HiSeq^™^ platform (2000 PE 100, Illumina, Inc., San Diego, CA, USA).

### 2.3 Bioinformatic Analyses

*De novo* assemblies were conducted on single samples to recover metagenome assembled genomes (MAGs). These methods are extensively detailed in a previously study (C. Yancey et al., 2022). Briefly, assemblies were completed using Megahit (Li et al., 2015), with differential coverage read mapping achieved through bowtie2 (Langmead and Salzberg, 2012). Multiple binning software were implemented to recover the highest quality MAGs and included Concoct (Alneberg et al., 2013), Metabat (Kang et al., 2015), Tetra-ESOM (Ultsch and Mörchen, 2009) and VizBin (Laczny et al., 2015). MAGs generated from all four binning software were run through DASTool (Sieber et al., 2018) to assess and choose the highest qualities MAGs for further analysis. For further MAG refinement, bins were manually assessed and curated using Anvi’o v.5 (Eren et al., 2015) and taxonomic confirmation, completion, strain heterogeneity, and contamination was further assessed with CheckM (Parks et al., 2015). From this effort, several cyanobacteria MAGs were generated including 9 *Microcystis* MAGs (C. Yancey et al., 2022), 2 ADA MAGs, and 6 *Pseudanabaena* MAGs (data not shown).

To understand the genetic context for the recovered ADA MAGs, the software pyani v.0.2.10 (Pritchard et al., 2015) was used to calculate average nucleotide identity between genomes. Calculations were completed on a large database of ADA genomes that included the two new ADA MAGs generated in this study as well as all the publicly available genomes for the ADA clade on NCBI (https://www.ncbi.nlm.nih.gov/, accessed April 2022). Pyani was run on default parameters togenerates pairwise ANI scores.

Relative abundance of ADA cyanobacteria was quantified as follows. Metagenomic reads were mapped to several cyanobacteria MAGs generated from the 2014 dataset. These MAGs included the two identified ADA MAGs, and a *Microcystis* MAG identified and examined in a previous study (C. Yancey et al., 2022), that were generated via *de novo* assembly from these samples. Reads mapped via Basic Alignment Search Tool (BLAST) (Madden, 2013) v.2.11.0 were kept for quantification if they satisfied a 95% identity and 80% alignment length cut off which was been previously shown to be sensitive and sufficient for metagenomic read mapping (C. E. Yancey et al., 2022). Reads per kilobase per million reads (RPKM) was calculated for each MAG to determine relative abundance using the equation below (Dick, 2018):

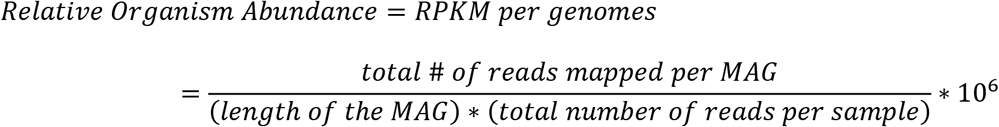

The software antiSMASH v.6.0 (Blin et al., 2021) was used to mine and annotate the ADA MAGs for biosynthetic gene clusters which encode secondary metabolites. Default parameters were run under the “relaxed” annotation settings. Identified BGCs were further deeply annotated on a by gene basis using blastP (Madden, 2013), and generated output from antiSMASH.

### 2.4 Relative Transcript Abundance

Relative abundance of transcripts was calculated as follows. Metatranscriptomic reads were mapped onto a database that contained all BGCs identified from the ADA MAGs as well as BGCs identified from *Microcystis* MAGs from a previous study (C. Yancey et al., 2022). This was done to ensure competitive mapping, especially for anabaenopeptin genes present in both the ADA and *Microcystis* genomes. Reads were mapped using BLAST and were kept if they had at least 95% identity and 80% alignment to the query. Singular top hits that met these criteria were then quantified. Similarly, a set of phosphorus and nitrogen genes were also subjected to metatranscriptomic read mapping under the same parameters listed above to determine nutrient uptake and metabolism throughout the bloom. The list of genes used for this analysis can be found of Table S1. Metatranscriptomic reads were also mapped to ADA MAGs using the same software and parameters listed above. This was used to calculate the relative transcript abundance for each BGC and nutrient gene using the equation below (Dick, 2018):

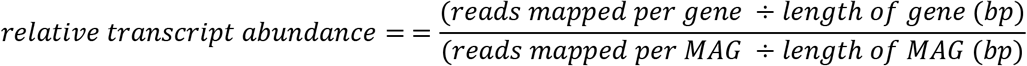

### 2.5 Figure Generation

Figures were generated using R and RStudio (Allaire, 2015) with the packages ggplot2 (Wickham, 2011), and ggpubr (Kassambara, 2020). Adobe Illustrator was used to render multiple paneled figures (“Adobe Illustrator,”2019).

## 3. Results

### 3.1 Overview of ADA cyanobacteria MAGs

From fifteen samples analyzed, d*e novo* metagenomic assembly produced two high quality ADA clade MAGs from two distinct samples (Fig 1A): station WE12 on August 4^th^, which was near the Toledo drinking water intake during the “do not drink” advisory and had the first observed peak of cyanobacterial biomass and peak of microcystins concentration (Berry et al., 2017); and WE4, an offshore station, which had low cyanobacterial biomass and concentration of microcystins on September 29^th^. These MAGs have high completion (>97%) and low contamination (<1%) as well as large N50s (>25,000 bp) (Fig 1B).

**Figure 1:**
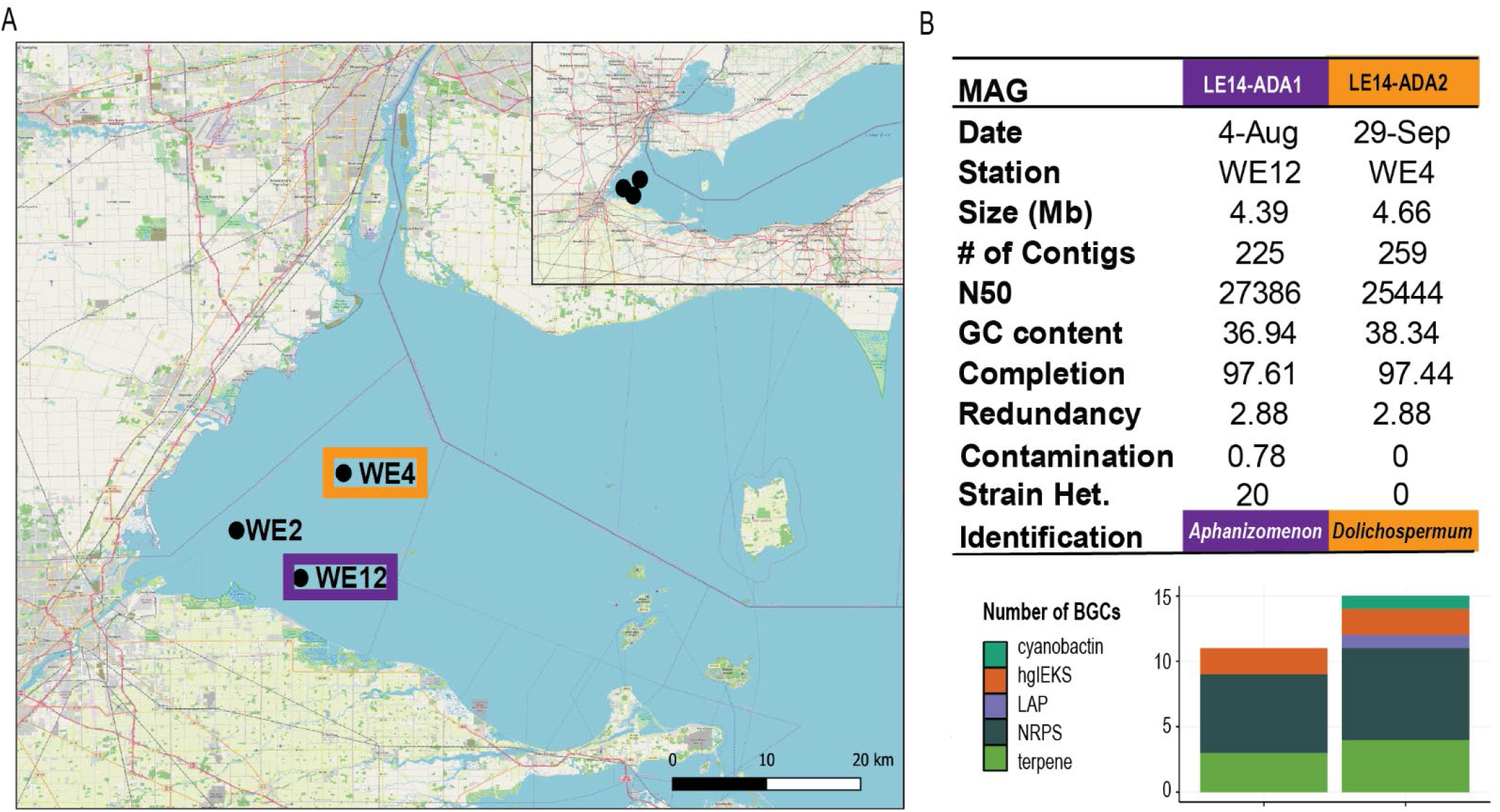
A) Map of Western Lake Erie and sampling stations in which ADA MAGs were recovered. LE14-ADA1 was recovered from WE12 (purple) and LE14-ADA2 was recovered from WE4 (orange). B) Summary statistics for ADA MAGs recovered from 2014 W. Lake Erie cyanoHAB, including sequences annotated as BGCs via antiSMASH v.6.0. Figure 1A was generated via QGIS using the Open Street Map (OSM) as a basemap (https://wiki.osmfoundation.org/wiki/Main_Page).

Analysis of pairwise average nucleotide identity (ANI) scores revealed that LE14-ADA 1 and LE14-ADA 2 had low ANI (88.8%) and clustered into distinct ADA clades, indicating that they are different species. LE14-ADA 1 clusters within a clade of nearly exclusively *Aphanizomenon* and is most closely related to *Aphanizomenon flos aquae* FACHB-1416 (ANI=0.989957, NCBI Genbank: GCA_014698695.1), which was isolated from China. LE14- ADA 2 belongs to a clade consisting mainly of *Dolichospermum* and is most closely related to a strain isolated from the western arm of Lake Superior, SB001 ((Sheik et al., 2022), ANI= 0.985575, NCBI BioSample: SAMN16655444). LE14-ADA 2 and SB001 fall within a clade that contains *Anabaena* sp. strains LE011-02 and AL09 (Driscoll et al., 2018) isolated from Lake Erie and Lake Ontario respectively (Fig. 2), thus representing a Great Lakes clade.

**Figure 2:**
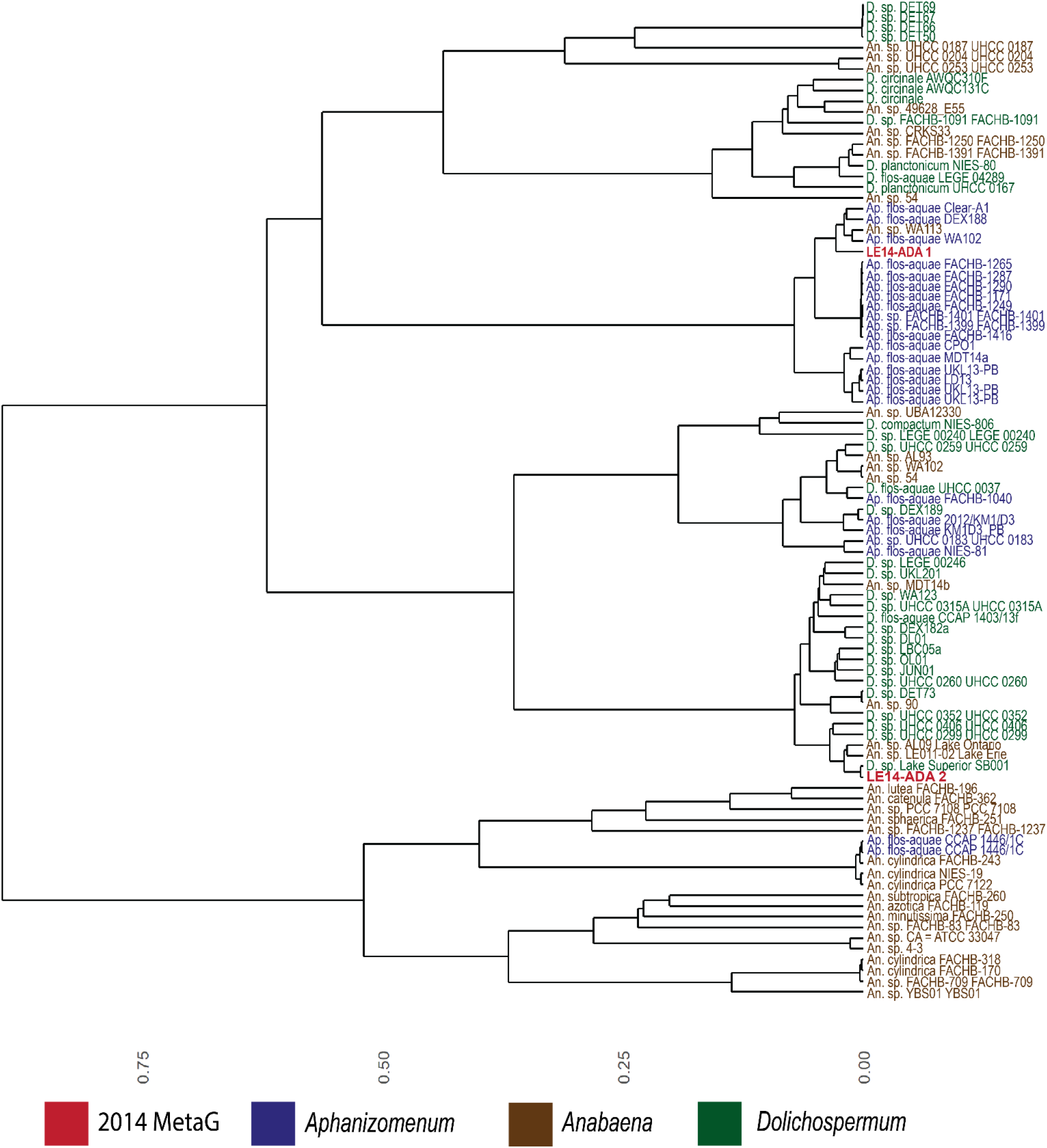
Cladogram of the ADA clade (includes all publicly available genomes from NCBI, accessed April 2022) based on average nucleotide identity between genomes. 2014 ADA MAGs LE14-ADA 1 and 2 are shown in red.

#### Temporal and Spatial Variation in Abundance of MAGs from ADA and Cyanobacteria

Mapping of metagenomic reads to MAGs provided estimates of ADA relative abundance across stations and sampling dates. LE14-ADA 1 reached maximum abundance when *Microcystis* was most abundant at nearshore stations August 4^th^ and September 29^th^ (Fig. 3A), and the relative abundances of ADA 1 and *Microcystis* were positively correlated (R=0.859, p=4.06e-05, Table S2). LE14-ADA 2 was less abundant throughout but its reached greatest abundance at offshore station WE4 on 29-Sep, when *Microcystis* abundance was low. There were no correlations between the relative abundance of LE14-ADA2 and *Microcystis* (Table S2). *Microcystis* spp. consistently dominated throughout the bloom and was 2-3 orders of magnitude more abundant than the ADA taxa (Fig 3A).

**Figure 3:**
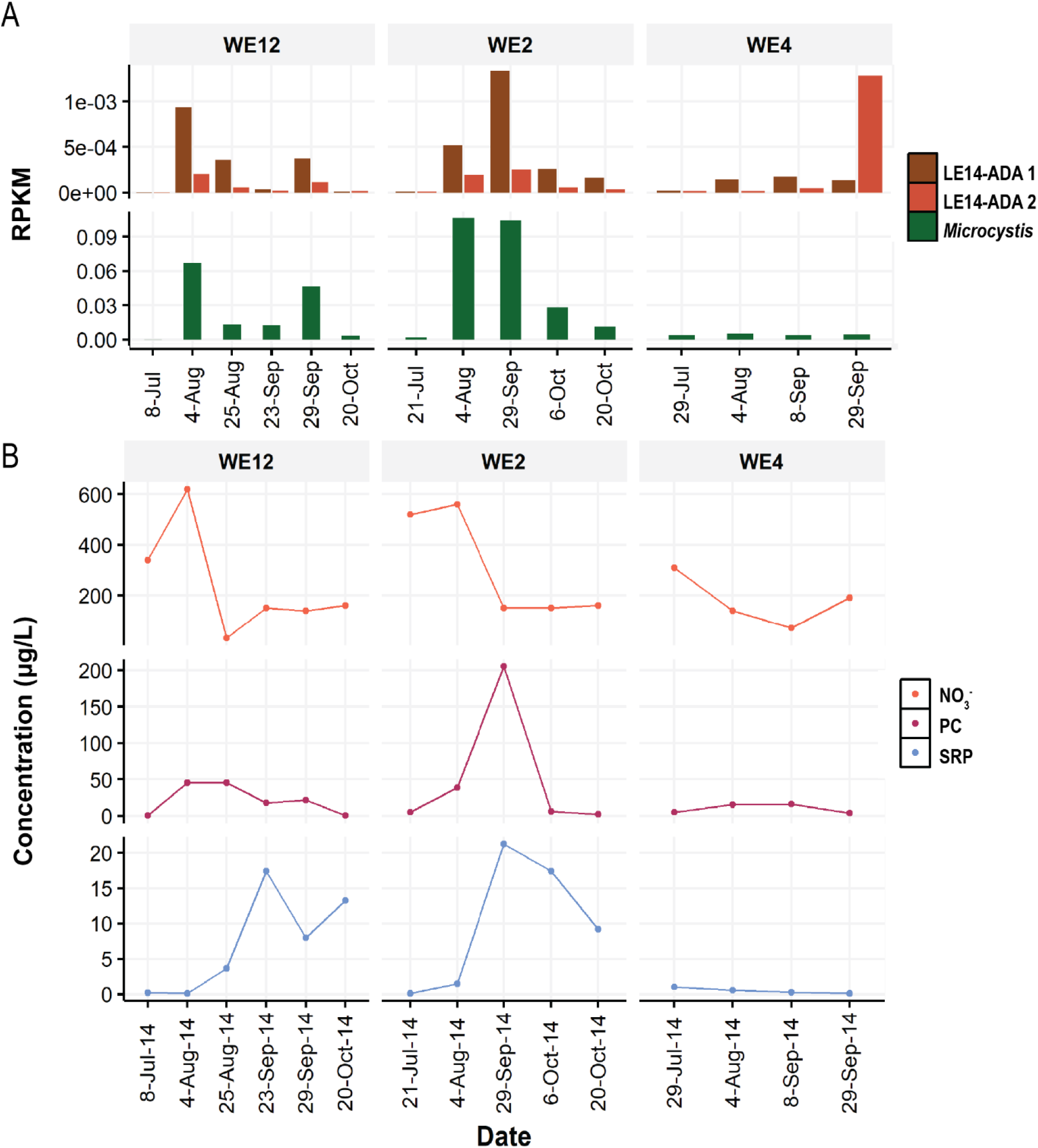
Relative abundance estimates of *Microcystis* and ADA as determined by mapping metagenomic reads onto MAGs for each taxa. Relative abundance was estimated via reads per kilobase million (RPKM, see methods). Note the difference in scale between *Microcystis* and ADA plots.

### 3.2 Biosynthetic Gene Clusters encoding secondary metabolites

While the *mcy* operon was not detected in either MAG, both contain over 10 biosynthetic gene clusters (BGCs) with nonribosomal peptide synthases (NRPS) and terpenes being the most common (Fig 1B). BGC mining via antiSMASH (Blin et al., 2021) also revealed the presence of 2 hgIE-KS clusters, which are believed to be important for heterocyst glycolipid formation (Fig 1B). LE14-ADA 2 contained more BGCs and a greater diversity of BGCs, including those encoding a cyanobactin and a linear azol(in)e-containing peptide (LAP) cluster (Fig 1B). Two clusters identified from LE14-ADA 2 were 100% similar (contained all genes to closest known BGC from the Minimum Information about a Biosynthetic Gene Cluster (MiBIG) database with significant BLAST hits) to previously described cyanopeptolin and geosmin encoding clusters (Fig. 4A, 4B). Cyanopeptolins inhibit several enzymes such as eukaryotic proteases including chymotrypsin (Bister et al., 2004; Gademann et al., 2010), while geosmin is a taste and odor compound (Gerber and Lechevalier, 1965; Izaguirre and Taylor, 2004). The genome of LE14- ADA 1 contained clusters with lower similarity hits to trypsin inhibitor psuedospumigin A/B/C/D/E/F ((Jokela et al., 2017), 66%), and protease inhibitor aeruginoside ((Ishida et al., 2007), 29%) suggesting they encode uncharacterized compounds related to identified metabolite classes (Fig 4C, 4D). Other BGCs with percent similarity hits to sequences in the MiBIG database are described in Table S3.

**Figure 4:**
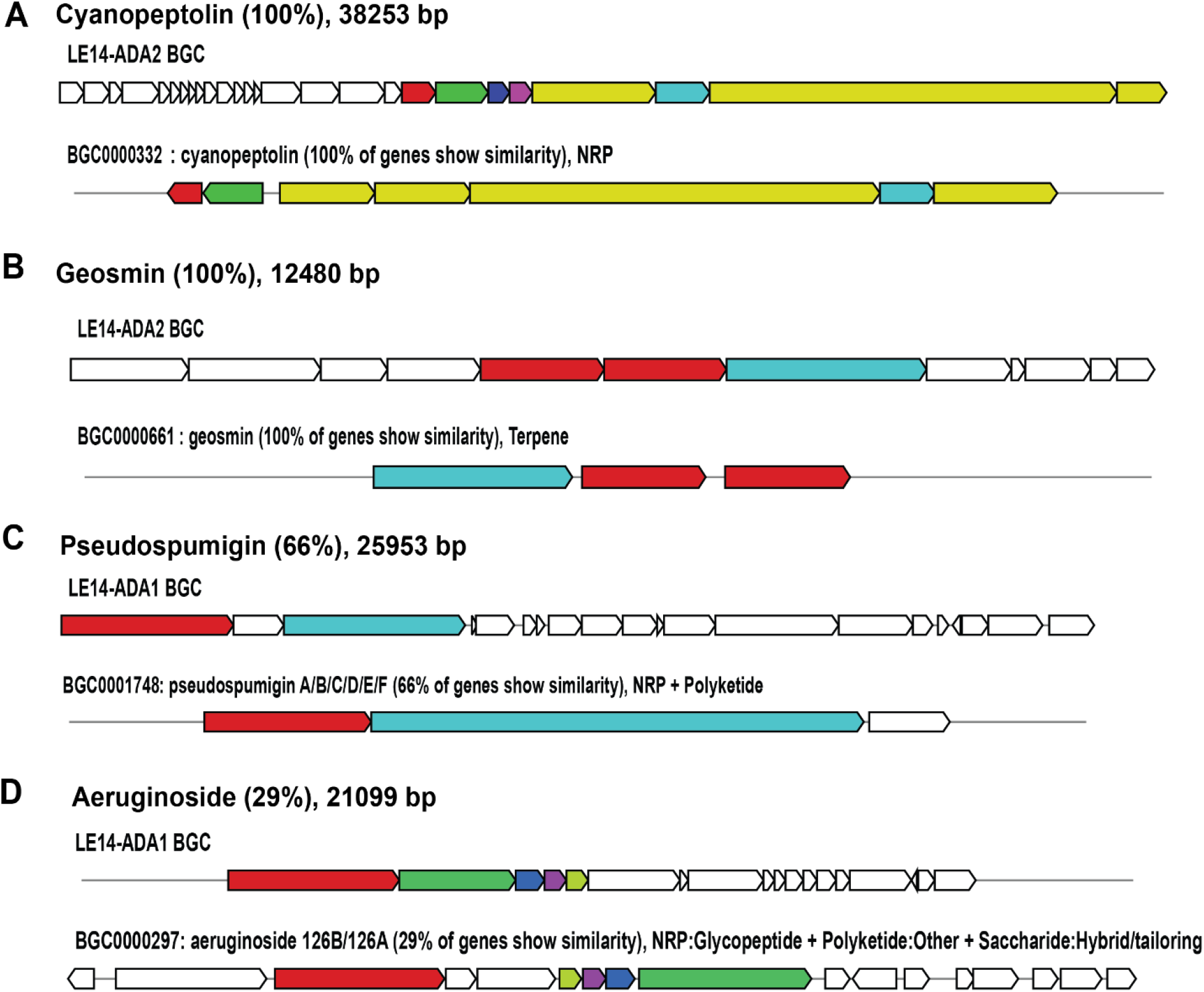
Schematics of selected BGCs as identified by antiSMASH v.6. The closest hit from the MiBIG repository is included for comparison. Schematics were generated as part of antiSMASH output.

### 3.3 Relative transcript abundance of ADA cyanobacterial genes

BGC expression, as estimated through relative transcript abundances, was greatest at nearshore stations and peak and late phases of the bloom for both ADA strains (Fig 5). There was no measurable expression during July at WE2 and WE4, when both ADA strains are low abundance. BGC expression generally increased during August, and late September-October (Fig 5). Some of the most highly expressed clusters include terpenes, which may encode taste and odor compounds or other novel secondary metabolites (Dittmann et al., 2015). The aeruginoside like cluster (29% similarity) from LE14-ADA 1 was one of the most highly expressed clusters from this strain both at nearshore and offshore stations (Fig 5A) while terpene 5 from LE14- ADA 2 was consistently expressed from peak to late phases at all three stations (Fig 5B).

**Figure 5:**
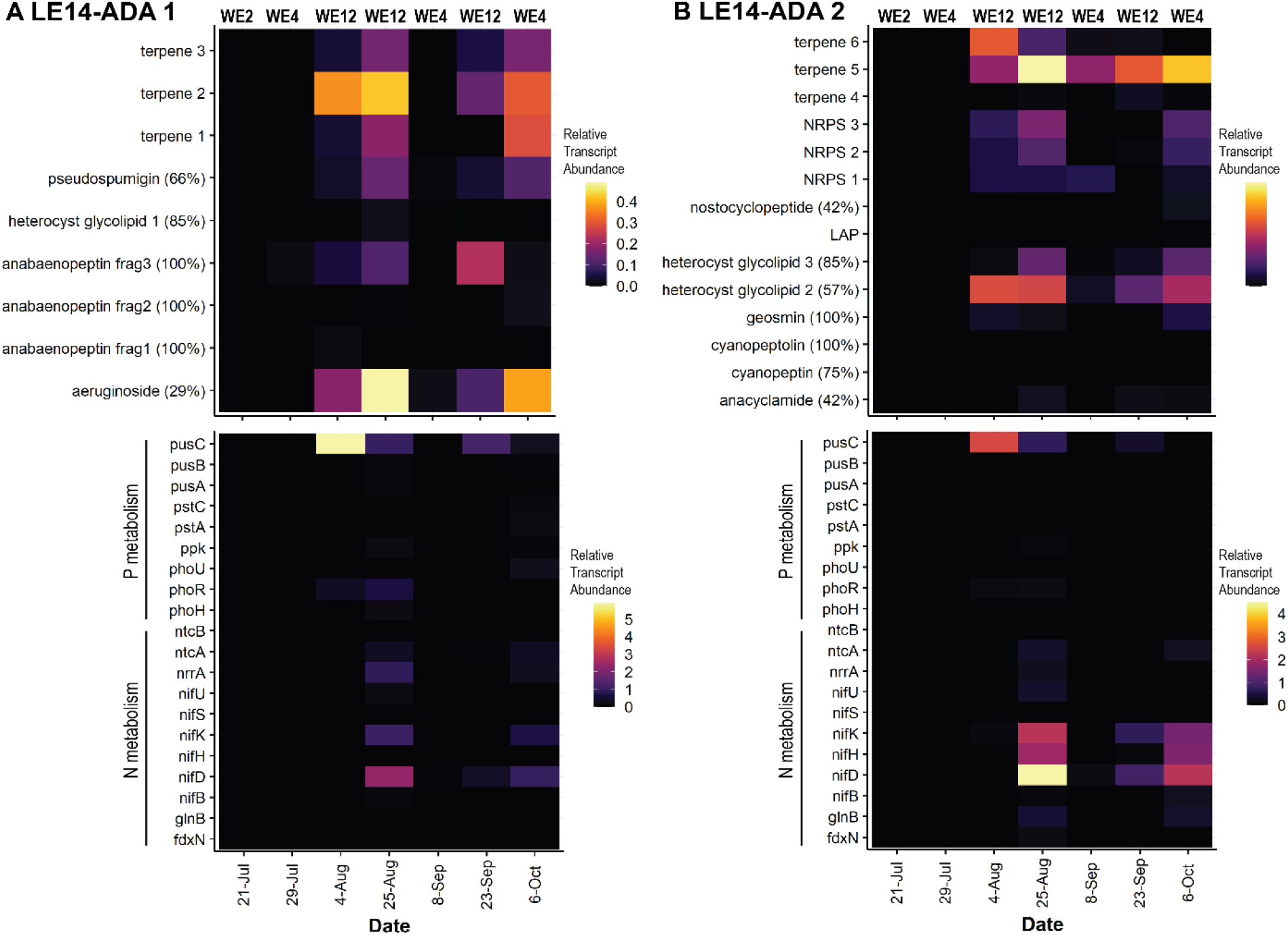
Relative transcript for both BGCs and nutrient uptake/metabolism genes from ADA MAGs. Expression estimates were calculated by mapping metatranscriptomic reads onto BGCs identified via antiSMASH. Percent similarities are listed next to BGCs with putative identifications. Nutrient metabolism gene transcript abundance is depicted in the bottom row. Relative transcript abundance calculations were completed for both A) LE14-ADA1 and B) LE14-ADA2.

Expression of genes for nutrient metabolism and uptake also varied between strains and throughout the bloom (Table 1, Fig. 5B). During August and late September, both ADA strains had greatest relative transcript abundance of *pusC,* an ABC phosphate transporter, suggesting these ADA strains were actively taking up exogenous phosphate from the water column. On 25 August, LE14-ADA 1 had higher transcript abundance of nitrogen fixation genes *nifD* and *nifH* (Fig. 5A). A similar pattern is observed in LE14-ADA 2; however, this trend persists throughout the rest of the bloom into early October. Higher relative transcript abundance of another N-fixation gene, *nifK* was also observed in LE14-ADA 2, as well as heterocyst glycolipid gene clusters which may aid in heterocyst formation ((Kampa et al., 2013), Fig. 5B). To assess ADA’s relative contribution to N-fixation in the broader cyanoHAB community, we searched metagenomic assemblies for nitrogen fixation genes belonging to other bacterial taxa, as it has become increasingly recognized that heterotrophs are capable of fixing N in aquatic systems (Bentzon-Tilia et al., 2015; Davis et al., 2015; Shiozaki et al., 2014). The only N-fixation genes found to be present in the 2014 cyanoHAB metagenome belonged to ADA clade of cyanobacteria, suggesting they are the primary, or sole, N-fixers within this system (Table S4).

**Table 1:**
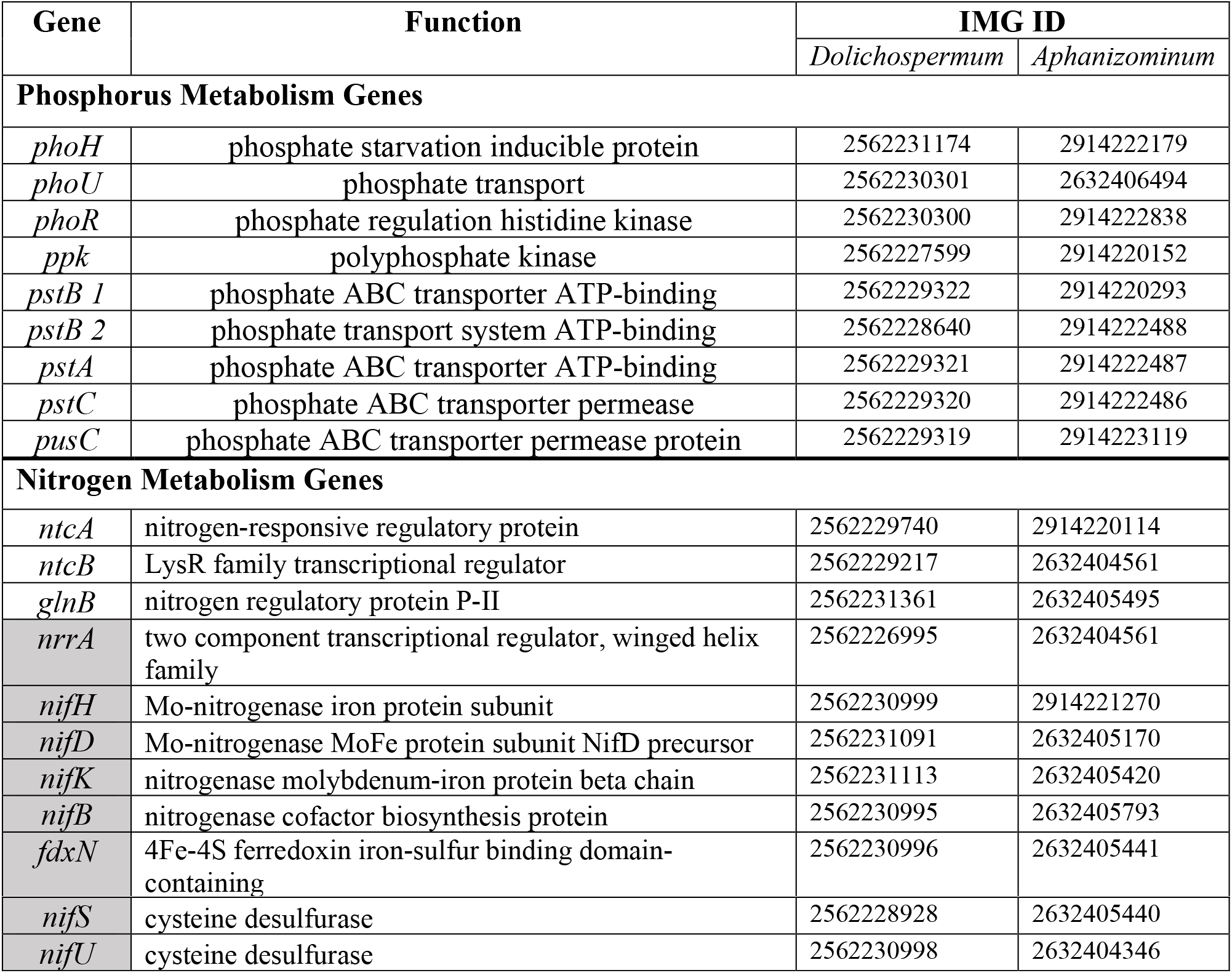
Functional roles of N and P genes. Boxes colored gray are genes involved in N fixation

## 4. Discussion

Metagenomics and metatranscriptomics provide detailed insights into the diversity of natural cyanoHAB communities, the ecophysiology and biosynthetic potential of the constituent species, and how they respond to environmental conditions. The recovery of highly complete MAGs and paired metatranscriptomic data presented an opportunity to assess the diversity and function and secondary metabolism of ADA cyanobacteria *in situ* through changing environmental conditions and community composition across the season in western Lake Erie. Although ADA cyanobacteria never dominated the 2014 cyanoHAB, our results suggest that they may play important roles in production of secondary metabolites and contributions to bioavailable N via N-fixation.

The ADA taxa were minor but pervasive members of the microbial community at all stations and times sampled during the 2014 western Lake Erie cyanoHAB, consistent with previous studies (Berry et al., 2017; Jankowiak et al., 2019; Steffen et al., 2017). Their relative abundance was dynamic, and the two ADA MAGs showed distinct spatiotemporal trends. These findings generally agree with previous studies (Berry et al., 2017; Jankowiak et al., 2019), although our approach implements metagenomic mapping onto entire MAGs instead of gene-targeted approaches. The *Aphanizomenon-*like MAG was associated with *Microcystis* blooms and most abundant at nearshore stations whereas the *Dolichospermum*-like LE14-ADA 2 MAG was most abundant offshore, late in the season, when *Microcystis* abundance was low. *Dolichospermum* spp. can also dominate Great Lakes cyanoHABs, for example preceding *Microcystis* as the dominate cyanobacterium early in blooms of the central basin of Lake Erie (Chaffin et al., 2019) or in Lake Superior (Sheik et al., 2022). *Aphanizomenon* spp. appear to be rarer now in the open waters of the Great Lakes but can dominate blooms in tributaries (McKay et al., 2020) despite dominating Lake Erie blooms in the 1960s and 1970s (Davis, 1964; Munawar and Munawar, 2011). This dynamic distribution of ADA organisms in time and space, taken together with evidence of potential for biosynthesis of saxitoxin (Chaffin et al., 2019) and guanitoxin (Lima et al., 2022) in the Great Lakes, underscore the need to determine environmental and ecological controls on their abundance.

According to a recently proposed phylogeny-based reorganization of the ADA clade into 10 species (Dreher et al., 2021), LE14-ADA 1 would be classified as “species 4”, and LE14-ADA 2 would be “species 2”, which both contain strains primarily from the United States (Dreher et al., 2021). Similarly, Österholm et al., 2020 divided the ADA group into seven distinct clades via phylogenomics. Based on this analysis, LE14-ADA 1 belongs in the Ò clade, all of which produce aeruginosins and are from freshwater sources in the United States. Genes that encode an aeruginoside-like compound, which is a class of aeruginosins, were found within the LE14-ADA 1 MAG. LE14-ADA 2 clusters most similarly with the α clade, which is comprised of strains from both Finland and the United States, with greater BGC diversity including genes that encode for microcystins (4/9 strains), and anabaenopeptins (8/9 strains) (Österholm et al., 2020). While a consensus on taxonomic classification of the ADA clade has yet to be reached, these results underscore the great genetic diversity of species within this group.

Each ADA MAG contains a distinct suite of transcriptionally active BGCs encoding secondary metabolites (Fig 1B). Two transcriptionally active BGCs from LE14-ADA 1 may encode toxic compounds, aeruginoside (29%) and psuedospumigin (66%) (Fig 5A). Aeruginosides belong to the peptide class aeruginosins and have been characterized in *Planktothrix* (Ishida et al., 2007). While the two core NRPS biosynthetic genes required for aeruginosin biosynthesis appear to be conserved (data not shown), the low percent similarity of LE14-ADA 1 aeruginoside cluster to the MiBIG database suggests that it may encode a novel aeruginosin or related compound. Likewise, a cluster predicted to encode pseudospumigin-like compound, a linear peptide related to aeruginosins and spumigins, may inhibit trypsin (Jokela et al., 2017).

Although low in transcriptional activity, LE14-ADA 2 contained several BGCs known to encode toxic compounds including related compounds cyanopeptolin (Tooming-Klunderud et al., 2007) and cyanopeptin (Rounge et al., 2007) as well as taste and odor compound geosmin (Gerber and Lechevalier, 1965). Several other BGCs with high transcriptional activity do not have known associated compounds, including the terpene and NRPS classes observed in both LE14-ADA 1 and 2. Similar trends are observed in *Microcystis* genomes isolated from the same bloom (C. Yancey et al., 2022). Together, this highlights our limited understanding of the breadth of cyanobacterial secondary metabolism within western Lake Erie and the need the assess potential risks of these secondary metabolites to ecosystem and human health.

The dynamic relative abundance of transcripts from nutrient uptake and metabolism genes of ADA cyanobacteria suggest shifts in nutrient acquisition strategies along with changing nutrient availability throughout the bloom. During low phosphate conditions of the peak bloom phase at WE12 (Fig. 3B), phosphate ABC transporter *pusC* was highly expressed by both ADA strains (Fig 5), suggesting P-limitation, stress, and competition with *Microcystis,* which is efficient at uptake and storage of inorganic P (Jacobson and Halmann, 1982).

We observed differential expression of various genes involved in N-fixation across time and space in western Lake Erie. This finding, along with previously reported expression of *nifH* (Steffen et al., 2015) and fixation of N in western Lake Erie (Natwora and Sheik, 2021), challenge the notion that current western Lake Erie cyanoHABs do not contain N-fixers (Barnard et al., 2021; Newell et al., 2019). Our results also suggest that the ADA taxa are the primary or sole N-fixers in western Lake Erie, in contrast to Sandusky Bay where N fixation has been attributed to both cyanobacteria and heterotrophic bacteria (Davis et al., 2015). Diazotrophy may be a critical strategy by which ADA cyanobacteria compete given that *Microcystis* efficiently scavenges N (Takamura et al., 1987; Wang et al., 2021); late season blooms are often limited by exogenous N (Chaffin et al., 2013), and secondary blooms of ADA cyanobacteria may be enabled by N fixation (Michalak et al., 2013). Alternatively, N fixation by ADA cyanobacteria may benefit *Microcystis* by mitigating N limitation. N rich amino acids and ammonium may “leak” from N-fixers and provide N to cyanoHABs (Ohlendieck et al., 2000; Wetzel, 2001), thereby providing resources for continued *Microcystis* dominance. Low abundance N-fixing cyanobacteria may serve as keystone species by fixing large quantities of N, even under conditions in which bioavailable N is abundant (Shiozaki et al., 2020). More work is needed address the impact of “leaky” N-fixation on Lake Erie cyanoHABs.

## 5. Conclusion

Other cyanobacteria present in western Lake Erie rarely receive attention due to the prevalence and threats posed by *Microcystis*. However, this study highlights two unique strains of ADA cyanobacteria that are pervasively present at low abundance and have the potential to produce a variety of known and unknown secondary metabolites. Their biosynthetic repertoire and transcriptional activity are varied and may reflect changes in bloom conditions as they compete for N and P and use alternate strategies for N acquisition and growth. N-fixation may help satisfy N demand and support the synthesis of N-rich secondary metabolites, especially during N deplete conditions, though the significance of N fixation for ADA organisms and the broader community remains to be quantified. While the monitoring of *Microcystis* and the production of its secondary metabolites is critical within western Lake Erie, this study demonstrates the need to further expand efforts toward other cyanobacteria found within these blooms as they may produce unmonitored toxins and shape observed community and ecosystem dynamics.

## Acknowledgements

We thank Dack Stuart and Kent Baker for field sampling support, and the NOAA Great Lakes Environmental Research Lab and Cooperative Institute for Great Lakes Research for allowing us to sample alongside their HAB monitoring efforts and providing seasonal nutrient data.

