## Supplemental Table for "Transcriptionally active nitrogen fixation and biosynthesis of diverse secondary metabolites by *Dolichospermum* and *Aphanizominom*-like Cyanobacteria in western Lake Erie Microcystis blooms"

**Supplemental Table 1**: P and N metabolism genes used for transcriptomic analysis

| **Phosphorus Genes** | | | | | | | | | | |
| --- | --- | --- | --- | --- | --- | --- | --- | --- | --- | --- |
|  | ***Anabaena* sp. 90** | | | | | ***Aphanizominum* FACHB 1287, 2012/KM/D3** | | | | |
| **Gene** | Gene ID | Locus Tag | Length | 42896 | 49633 | Gene ID | Locus Tag | Length | 42896 | 49633 |
| phoH | [2562231174](https://img.jgi.doe.gov/cgi-bin/mer/main.cgi?section=GeneDetail&page=geneDetail&gene_oid=2562231174) | ANA_C13591 | 954 | Y | Y | [2914222179](https://img.jgi.doe.gov/cgi-bin/mer/main.cgi?section=GeneDetail&page=geneDetail&gene_oid=2914222179) | Ga0477648_45_138477_139430 | 954 | Y | Y |
| phoU | [2562230301](https://img.jgi.doe.gov/cgi-bin/mer/main.cgi?section=GeneDetail&page=geneDetail&gene_oid=2562230301) | ANA_C12729 | 669 | Y | Y | [2632406494](https://img.jgi.doe.gov/cgi-bin/mer/main.cgi?section=GeneDetail&page=geneDetail&gene_oid=2632406494) | Ga0070878_11848 | 390 | Y | Y |
| phoR | [2562230300](https://img.jgi.doe.gov/cgi-bin/mer/main.cgi?section=GeneDetail&page=geneDetail&gene_oid=2562230300) | ANA_C12728 | 1320 | Y | Y | [2914222838](https://img.jgi.doe.gov/cgi-bin/mer/main.cgi?section=GeneDetail&page=geneDetail&gene_oid=2914222838) | Ga0477648_69_6114_7421 | 1308 | Y | Y |
| ppk | [2562227599](https://img.jgi.doe.gov/cgi-bin/mer/main.cgi?section=GeneDetail&page=geneDetail&gene_oid=2562227599) | ANA_C10060 | 2166 | Y | Y | [2914220152](https://img.jgi.doe.gov/cgi-bin/mer/main.cgi?section=GeneDetail&page=geneDetail&gene_oid=2914220152) | Ga0477648_12_22313_24478 | 2166 | Y | Y |
| pusA | [2562229322](https://img.jgi.doe.gov/cgi-bin/mer/main.cgi?section=GeneDetail&page=geneDetail&gene_oid=2562229322) | ANA_C11758 | 780 | Y | Y | [2914220293](https://img.jgi.doe.gov/cgi-bin/mer/main.cgi?section=GeneDetail&page=geneDetail&gene_oid=2914220293) | Ga0477648_12_152706_153488 | 783 | Y | Y |
| pusB | [2562228640](https://img.jgi.doe.gov/cgi-bin/mer/main.cgi?section=GeneDetail&page=geneDetail&gene_oid=2562228640) | ANA_C11088 | 699 |  | Y | [2914222488](https://img.jgi.doe.gov/cgi-bin/mer/main.cgi?section=GeneDetail&page=geneDetail&gene_oid=2914222488) | Ga0477648_56_100694_101500 | 807 | Y | Y |
| pstA | 2562229321 | ANA_C11757 | 873 | Y | Y | [2914222487](https://img.jgi.doe.gov/cgi-bin/mer/main.cgi?section=GeneDetail&page=geneDetail&gene_oid=2914222487) | Ga0477648_56_99755_100636 | 882 | Y | Y |
| pstC | 2562229320 | ANA_C11756 | 999 | Y | Y | [2914222486](https://img.jgi.doe.gov/cgi-bin/mer/main.cgi?section=GeneDetail&page=geneDetail&gene_oid=2914222486) | Ga0477648_56_98766_99710 | 945 | Y | Y |
| pusC | 2562229319 | ANA_C11755 | 1014 | Y | Y | 2914223119 | Ga0477648_85_79251_80291 | 1041 | Y | Y |
| **Nitrogen Genes** | | | | | | | | | | |
|  | ***Anabaena* sp. 90** | | | | | ***Aphanizominum* FACHB 1287, 2012/KM/D3** | | | | |
| **Gene** | Gene ID | Locus Tag | Length | 42896 | 49633 | Gene ID | Locus Tag | Length | 42896 | 49633 |
| ntcA | [2562229740](https://img.jgi.doe.gov/cgi-bin/mer/main.cgi?section=GeneDetail&page=geneDetail&gene_oid=2562229740) | ANA_C12173 | 672 | Y | Y | [2914220114](https://img.jgi.doe.gov/cgi-bin/mer/main.cgi?section=GeneDetail&page=geneDetail&gene_oid=2914220114) | Ga0477648_11_52532_53203 | 672 | Y | Y |
| glnB | [2562229217](https://img.jgi.doe.gov/cgi-bin/mer/main.cgi?section=GeneDetail&page=geneDetail&gene_oid=2562229217) | ANA_C11655 | 339 | Y | Y | 2632405495 | Ga0070878_113511 | 339 | Y | Y |
| ntcB | [2562231361](https://img.jgi.doe.gov/cgi-bin/mer/main.cgi?section=GeneDetail&page=geneDetail&gene_oid=2562231361) | ANA_C13773 | 924 | Y | Y | 2632404561 | Ga0070878_108128 | 930 | Y | Y |
| nrrA | [2562226995](https://img.jgi.doe.gov/cgi-bin/mer/main.cgi?section=GeneDetail&page=geneDetail&gene_oid=2562226995) | ANA_C20328 |  | Y | Y | 2914221270 | Ga0477648_25_40471_41241 | 771 |  |  |
| nifH | [2562230999](https://img.jgi.doe.gov/cgi-bin/mer/main.cgi?section=GeneDetail&page=geneDetail&gene_oid=2562230999) | ANA_C13419 | 870 | Part | Part | [2632405170](https://img.jgi.doe.gov/cgi-bin/mer/main.cgi?section=GeneDetail&page=geneDetail&gene_oid=2914219458) | Ga0070878_111820 | 981 | Y | Y |
| nifD | [2562231091](https://img.jgi.doe.gov/cgi-bin/mer/main.cgi?section=GeneDetail&page=geneDetail&gene_oid=2562231091) | ANA_C13511 | 1503 | Y | Y | [2632405420](https://img.jgi.doe.gov/cgi-bin/mer/main.cgi?section=GeneDetail&page=geneDetail&gene_oid=2632405420) | Ga0070878_113212 | 1395 | Y | Y |
| nifK | [2562231113](https://img.jgi.doe.gov/cgi-bin/mer/main.cgi?section=GeneDetail&page=geneDetail&gene_oid=2562231113) | ANA_C13533 | 1536 | Y | Y | [2632405793](https://img.jgi.doe.gov/cgi-bin/mer/main.cgi?section=GeneDetail&page=geneDetail&gene_oid=2632405793) | Ga0070878_115018 | 1536 | Y | Y |
| nifB | [2562230995](https://img.jgi.doe.gov/cgi-bin/mer/main.cgi?section=GeneDetail&page=geneDetail&gene_oid=2562230995) | ANA_C13415 | 1440 | Part | Y | [2632405441](https://img.jgi.doe.gov/cgi-bin/mer/main.cgi?section=GeneDetail&page=geneDetail&gene_oid=2632405441) | Ga0070878_113233 | 1440 | Y | Y |
| fdxN | 2562230996 | ANA_C13416 | 345 | Y | Y | 2632405440 | Ga0070878_113232 | 345 | Y | Y |
| nifS | [2562228928](https://img.jgi.doe.gov/cgi-bin/mer/main.cgi?section=GeneDetail&page=geneDetail&gene_oid=2562228928) | ANA_C11370 | 1149 | Y | Y | 2632404346 | Ga0070878_107125 | 1149 | Y | Y |
| nifU | [2562230998](https://img.jgi.doe.gov/cgi-bin/mer/main.cgi?section=GeneDetail&page=geneDetail&gene_oid=2562230998) | ANA_C13418 | 900 | Y | Y | [2632405438](https://img.jgi.doe.gov/cgi-bin/mer/main.cgi?section=GeneDetail&page=geneDetail&gene_oid=2632405438) | Ga0070878_113230 | 900 | Y | Y |

**Supplemental Table 2**: Pearson’s correlations between the relative abundance of ADA cyanobacteria and *Microcystis*

| **Western Lake Erie MAG** | **R** | **p-value** |
| --- | --- | --- |
| LE14-ADA1 | 0.859 | 4.06e-05 |
| LE14-ADA2 | 0.067 | 0.811 |

**Supplemental Table 3**: Selected BGCs and their putative identification via antiSMASH v.6.0.

| **Genome** | **Contig_ID** | **Length** | **BGC_type** | **Put_ID** | **Similarity** |
| --- | --- | --- | --- | --- | --- |
| LE14-ADA1 | k119_242196 | 4300 | NRPS | anabaenopeptin NZ 857/nostamide A | 100 |
|  | k119_219210 | 4059 | NRPS | anabaenopeptin NZ 857/nostamide A | 100 |
|  | k119_93841 | 6158 | NRPS | anabaenopeptin NZ 857/nostamide A | 100 |
|  | k119_291716 | 12732 | hgIEKS | heterocyst glycolipids | 85 |
|  | k119_253808 | 30602 | NRPS | psuedospumigen A/B/C/D/E/F | 66 |
|  | k119_142266 | 27155 | NRPS | aeruginoside | 29 |
|  | k119_77657 | 23000 | hgIEKS | 6,6'-oxybis(2,4-dibromophenol) | 14 |
| LE14-ADA2 | k141_342326 | 15853 | terpene | geosmin | 100 |
|  | k141_557282 | 42095 | NRPS | cyanopeptolin | 100 |
|  | k141_295936 | 21236 | NRPS | cyanopeptolin | 57 |
|  | k141_97816 | 16866 | cyanobactin | anacyclamide | 50 |
|  | k141_327140 | 35361 | NRPS | nostocyclopeptide | 42 |
|  | k141_139283 | 14686 | NRPS | nostocyclopeptide | 37 |

**Supplemental Table 4:** BLAST for N fixation Genes. Contigs with N fixation genes found in metagenomic assemblies were BLASTed against the nonredundant NCBI database to determine taxonomic assignment.

| **Sample_ID** | **contig ID** | **gene** | **%ID** | **Alignment Length** | **Top NCBI Hit** | **NCBI Accession** | **NCBI %ID** |
| --- | --- | --- | --- | --- | --- | --- | --- |
| 42895 | No HITS | | | | | | |
| 42896 | 76095 | nifU | 95 | 900 | *Aphanizomenon flos-aquae* DEX188 chromosome, complete genome | CP051188.1 | 99 |
|  | 76095 | nifB | 93 | 924 | *Aphanizomenon flos-aquae* DEX188 chromosome, complete genome | CP051188.1 | 99 |
|  | 76095 | fdxN | 92 | 345 | *Aphanizomenon flos-aquae* DEX188 chromosome, complete genome | CP051188.1 | 99 |
|  | 126771 | nifD | 96 | 1382 | *Aphanizomenon flos-aquae* DEX188 chromosome, complete genome | CP051188.1 | 98.6 |
|  | 230629 | nifS | 86 | 1144 | *Aphanizomenon flos-aquae* DEX188 chromosome, complete genome | CP051188.1 | 99.6 |
|  | 242374 | nifB | 91 | 504 | *Aphanizomenon flos-aquae* DEX188 chromosome, complete genome | CP051188.1 | 99.7 |
|  | 244836 | nifK | 92 | 1536 | *Aphanizomenon flos-aquae* DEX188 chromosome, complete genome | CP051188.1 | 99 |
| 49615 | No HITS | | | | | | |
| 49622 | No HITS | | | | | | |
| 49625 | 4813 | nifD | 95 | 1382 | *Aphanizomenon flos-aquae* DEX188 chromosome, complete genome | CP051188.1 | 99 |
|  | 4813 | nifB | 91 | 1440 | *Aphanizomenon flos-aquae* DEX188 chromosome, complete genome | CP051188.1 | 99 |
|  | 4813 | nifU | 95 | 900 | *Aphanizomenon flos-aquae* DEX188 chromosome, complete genome | CP051188.1 | 99 |
|  | 4813 | fdxN | 92 | 345 | *Aphanizomenon flos-aquae* DEX188 chromosome, complete genome | CP051188.1 | 99 |
|  | 18623 | nifS | 86 | 1144 | *Aphanizomenon flos-aquae* DEX188 chromosome, complete genome | CP051188.1 | 99 |
|  | 19180 | nifK | 93 | 407 | *Aphanizomenon flos-aquae* DEX188 chromosome, complete genome | CP051188.2 | 95 |
|  | 20534 | nifU | 94 | 234 | *Dolichospermum heterosporum* TAC447 strain NIES-1697 chromosome, complete genome | CP099464.1 | 96 |
|  | 25640 | nifD | 93 | 152 | *Dolichospermum flos-aquae* CCAP 1403/13F chromosome, complete genome | CP051206.1 | 93 |
|  | 135622 | nifK | 94 | 310 | *Dolichospermum sp. DL01* chromosome, complete genome | CP050884.1 | 97 |
|  | 150129 | nifB | 93 | 350 | *Aphanizomenon flos-aquae* DEX188 chromosome, complete genome | CP051188.1 | 96 |
|  | 170665 | nifK | 92 | 1536 | *Aphanizomenon flos-aquae* DEX188 chromosome, complete genome | CP051188.2 | 98 |
| 49629 | 132092 | fdxN | 92 | 270 | *Aphanizomenon flos-aquae* DEX188 chromosome, complete genome | CP051188.3 | 94 |
|  | 148359 | nifK | 93 | 315 | *Aphanizomenon flos-aquae* DEX188 chromosome, complete genome | CP051188.4 | 96 |
|  | 175598 | nifU | 94 | 766 | *Aphanizomenon flos-aquae* DEX188 chromosome, complete genome | CP051188.5 | 95 |
|  | 230262 | nifS | 84 | 300 | *Dolichospermum* sp. DET69 chromosome, complete genome | CP070233.1 | 89 |
|  | 250934 | nifK | 93 | 955 | *Aphanizomenon flos-aquae* DEX188 chromosome, complete genome | CP051188.1 | 91 |
| 49633 | 740599 | nifK | 95 | 1536 | *Dolichospermum* sp. UKL201 chromosome, complete genome | CP050891.1 | 99 |
|  | 740599 | nifB | 96 | 1440 | *Dolichospermum* sp. UKL201 chromosome, complete genome | CP050891.1 | 99 |
|  | 740599 | nifD | 95 | 1394 | *Dolichospermum* sp. UKL201 chromosome, complete genome | CP050891.1 | 99 |
|  | 740599 | nifU | 95 | 900 | *Dolichospermum* sp. UKL201 chromosome, complete genome | CP050891.1 | 99 |
|  | 740599 | fdxN | 94 | 345 | *Dolichospermum* sp. UKL201 chromosome, complete genome | CP050891.1 | 99 |
|  | 788363 | nifS | 94 | 1150 | *Dolichospermum* sp. LBC05a chromosome, complete genome | CP050882.1 | 96 |
| 49636 | No HITS | | | | | | |
| 49637 | No HITS | | | | | | |
| 53598 | No HITS | | | | | | |
| 53599 | No HITS | | | | | | |
| 53600 | 21958 | nifS | 87 | 382 | *Aphanizomenon flos-aquae* DEX188 chromosome, complete genome | CP051188.1 | 98 |
|  | 55860 | nifD | 96 | 880 | *Aphanizomenon flos-aquae* DEX188 chromosome, complete genome | CP051188.2 | 100 |
|  | 68535 | nifU | 96 | 340 | *Aphanizomenon flos-aquae* DEX188 chromosome, complete genome | CP051188.2 | 99 |
|  | 79224 | nifB | 90 | 359 | *Aphanizomenon flos-aquae* DEX188 chromosome, complete genome | CP051188.3 | 100 |
|  | 212327 | nifS | 85 | 333 | *Aphanizomenon flos-aquae* DEX188 chromosome, complete genome | CP051188.4 | 99 |
|  | 238880 | nifK | 93 | 539 | *Aphanizomenon flos-aquae* DEX188 chromosome, complete genome | CP051188.5 | 100 |
|  | 298821 | nifK | 91 | 930 | *Aphanizomenon flos-aquae* DEX188 chromosome, complete genome | CP051188.6 | 99 |
|  | 306979 | nifD | 96 | 321 | *Aphanizomenon flos-aquae* DEX188 chromosome, complete genome | CP051188.7 | 100 |
| 53601 | 29863 | nifB | 90 | 332 | *Aphanizomenon flos-aquae* DEX188 chromosome, complete genome | CP051188.8 | 100 |
|  | 191267 | nifD | 94 | 365 | *Aphanizomenon flos-aquae* DEX188 chromosome, complete genome | CP051188.9 | 100 |
|  | 226050 | nifD | 97 | 390 | *Aphanizomenon flos-aquae* DEX188 chromosome, complete genome | CP051188.10 | 100 |
|  | 233917 | nifK | 92 | 399 | *Aphanizomenon flos-aquae* DEX188 chromosome, complete genome | CP051188.11 | 99 |
|  | 299651 | nifS | 86 | 838 | *Aphanizomenon flos-aquae* DEX188 chromosome, complete genome | CP051188.12 | 98 |
|  | 346083 | nifK | 92 | 298 | *Aphanizomenon flos-aquae* DEX188 chromosome, complete genome | CP051188.13 | 99 |
|  | 354690 | fdxN | 92 | 330 | *Aphanizomenon flos-aquae* DEX188 chromosome, complete genome | CP051188.14 | 100 |
|  | 360023 | nifD | 97 | 309 | *Aphanizomenon flos-aquae* DEX188 chromosome, complete genome | CP051188.15 | 100 |
| 53602 | No HITS | | | | | | |
| 53603 | 65119 | nifU | 95 | 351 | *Aphanizomenon flos-aquae* DEX188 chromosome, complete genome | CP051188.15 | 100 |
|  | 99205 | nifK | 93 | 449 | *Aphanizomenon flos-aquae* DEX188 chromosome, complete genome | CP051188.16 | 98 |
|  | 151042 | nifD | 95 | 1025 | *Aphanizomenon flos-aquae* DEX188 chromosome, complete genome | CP051188.17 | 98 |
|  | 178298 | nifK | 92 | 553 | *Aphanizomenon flos-aquae* DEX188 chromosome, complete genome | CP051188.18 | 98 |
|  | 207680 | nifB | 89 | 237 | *Aphanizomenon flos-aquae* DEX188 chromosome, complete genome | CP051188.19 | 99 |
